# VARIATION IN GAME AND DOMESTIC ANIMAL RATIOS IN THE 7TH - 5TH MILLENNIA BCE IN THE LOWER VOLGA REGION

**DOI:** 10.1101/2025.03.02.640075

**Authors:** A.A. Vybornov, H.C. Doga, P.A. Kosintsev, N.V. Roslyakova, P.F. Kuznetsov

## Abstract

This paper presents the results of analysis of the species composition found in sites dating to the 7th - 5th millennia BCE. These sites are either monocultural, or multicultural where the cultural layers belonging to different periods are separated from each other by sterile layers. As a result, we were able to trace the variation through time in the ratios of game/domestic animals in the Neolithic - Eneolithic periods. In the Early Neolithic, the kulan was the main game animal. During the Middle and Late Neolithic, hunting was diversified and such animals as saiga, aurochs, and horse, along with kulan, became the main target species. In the Early Eneolithic, the first domestic animals, i.e. sheep and goats, appeared. Cattle appeared in the Late Eneolithic. The share of game animals during this period sharply decreased, even to the point of the complete disappearance of such species as aurochs and horse.

## Introduction

The Stone Age sites in the Lower Volga and the North Caspian regions have been periodically studied over the past forty years. Since 2013, the archaeological expedition of the Samara State University of Social Sciences and Education has been excavating these settlements and temporary campsites on a yearly basis. The basic purpose of these studies is to search for and excavate the monocultural settlements that belong to one particular archaeological culture, or the multicultural settlements with well-defined cultural layers separated by sterile layers from each other. This research makes it possible to establish the actual chronology of the cultures and determine the composition of the meat diet of the ancient inhabitants. Currently, 11 single-layer and three multilayer settlements with interlaid sterile layers have been studied (Fig. 1). Previously the species composition of the animals hunted and kept by the Eastern European population has been determined based on refuse material obtained from the sites where the cultural layers of the Stone and Bronze Ages, the Early Iron Age and even the Middle Ages are not explicitly differentiated. Consequently, a novel objective to analyze the changes in the ratios of game animals in the course of two thousand years has been set and a solution has been suggested. It has also been found that domestic animals appeared in the Early Eneolithic. The horse remained a game animal during most of the period under review. With the appearance of domestic species in the Eneolithic, the proportion of horse remains sharply decreased, to the point of its complete disappearance.

**Fig. 1.**
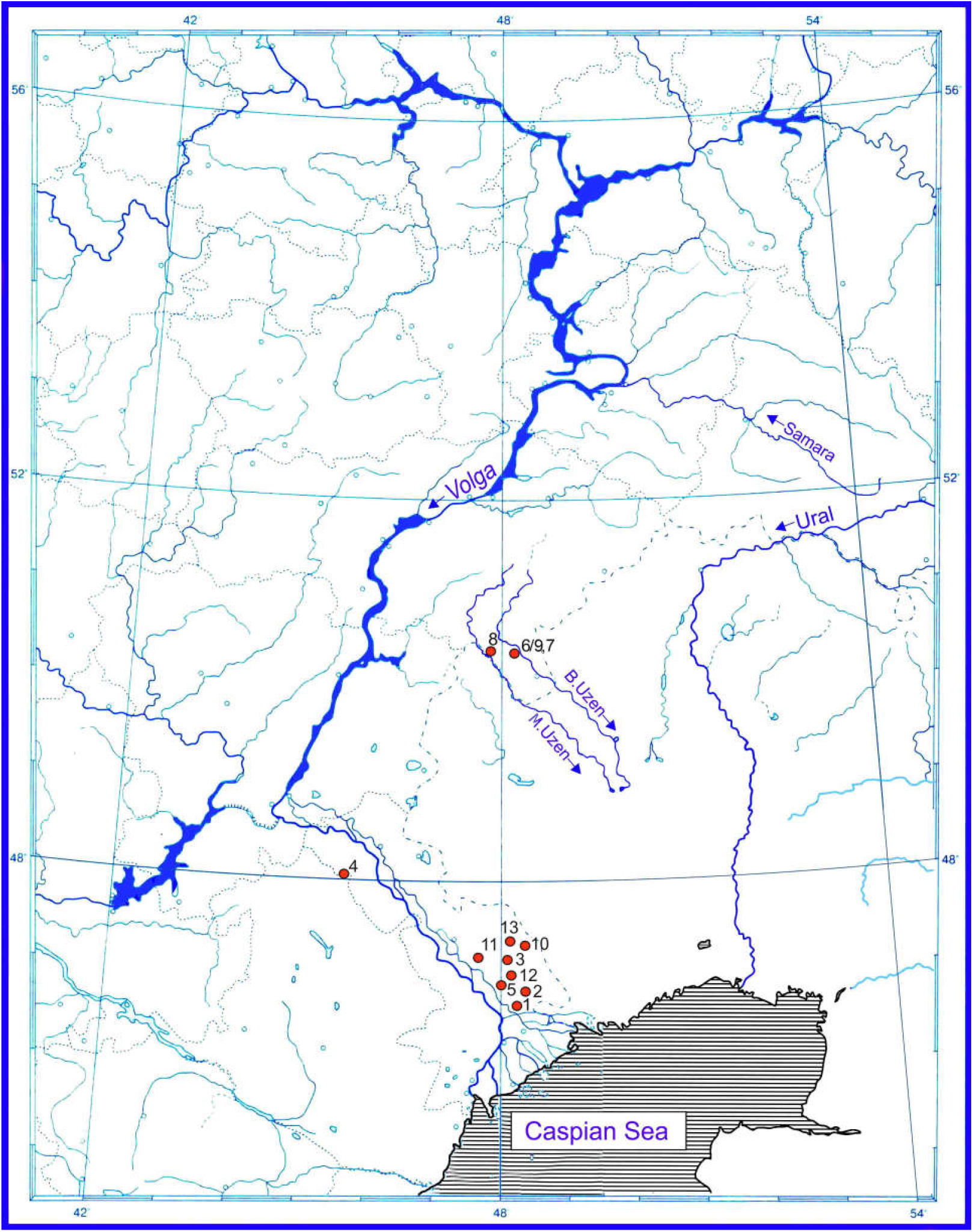
Map of the Lower Volga sites dating to the 7th-5th millennium BCE. 1-Baibek; 2-Kair-Shak III; 3-Tenteksor; 4-Jangar; 5-Priozernaya; 6-Oroshaemoe bottom layer; 7-Algay; 8-Varfolomeevskaya; 9-Oroshaemoe middle layer; 10-Kurpezhe-Molla; 11-Kara-Khuduk I; 12-Kair-Shak VI; 13-Kombak-Te.

### 1. Geographical characteristics of the region

The Lower Volga is represented by three landscapes: the North Caspian semi-desert (Astrakhan region), the southern steppes of the North-West Caspian (Kalmykia) and the Volga steppes (Volgograd and Saratov regions). According to paleogeographers, the natural environment and climate of this territory today do not differ much from the Neolithic and Eneolithic periods. Thus, in the Neolithic North Caspian area goosefoot (Chenopodiaceae) and wormwood (Artemisia) predominated, which is indicative of an arid climate. At a later period, the region becomes more humid, which is typical, except for short periods of aridity, of the Middle Eneolithic (Lavrushin et al., 1998). In the Volga steppes there is a change from semi-desert conditions in the Early Neolithic to typical steppe climate in the later Neolithic period, though in the Late Neolithic the climate becomes semi-desert again. In the Early Eneolithic period, apart from goosefoot and wormwood, mixed herbs and forbs are common for the region. The beginning of the Early Eneolithic is associated with some desiccation. Then humidity, which dominates in the Middle Eneolithic, increases (Ovchinnikov et al., 2022; Vybornov et al., 2022a). In the North-West Caspian region, the Middle Neolithic period was arid. This is confirmed by the calcareous layer in the second layer of the Jangar settlement. At a later period (the first Jangar layer), the climate changes and it becomes more humid (Koltsov, 2005).

### 2. Characteristics of the archaeological sites (Fig.1. Map)

The Neolithic and Eneolithic sites of the North Caspian are situated around the “sors” (prehistoric lakes) formed by the flooded areas of the Caspian Sea. At that time they were fresh-water lakes. These sites are similar to the North-West Caspian campsites located in the Sarpa Lakes area. In the Volga steppes the sites appeared and developed along the medium-sized rivers such as the Bolshoy Uzen and the Maly Uzen. In all the above cases, there is evidence for the ancient people being engaged in fishing (Grechkina et al., 2014; Vybornov et al., 2021a). Dwelling construction at the sites begins in the Middle Neolithic. In some settlements a number of dwellings apparently built at different times were found. This indicates a temporary (probably in winter) settled way of life of hunters and fishermen. This is also confirmed by the alternation between layers with cultural remains and sterile layers at a number of campsites of both the Neolithic and the Eneolithic periods.

The sites of the Middle Neolithic period in the North Caspian belong to the Kair-shak type of the Seroglazovo archaeological culture (the campsites of Kair-Shak III, Baibek, etc.). Pottery is made from lake silt. The pots are flat-bottomed. They are decorated with incisions and scattered impressions made by a pointed tool. The patterns are of geometric design. Stone tools, apart from scrapers and points, also include geometric microliths in the form of a segment. The campsites of the Late Neolithic (Tenteksor, Taskuduk, Priozernaya, etc.) belong to the Tenteksor culture. The pottery decoration of the period is characterized by impressions made with an obliquely held point. The flint assemblages include trapezes with a planed-off surface, as well as polished and drilled artifacts (Vybornov et al., 2020a). The campsites of the North-West Caspian region belong to the Jangar culture. The pottery is made from silt and silty clay. Flat-bottomed vessels are decorated with impressions made with an obliquely held point and incisions. The patterns are geometric. The majority of the stone tools include scrapers, points, geometric microliths: trapezes and segments (Koltsov, 2005). The Volga steppes are mainly represented by the Orlovka culture (Varfolomeevskaya camp, Orlovka, Algay, etc.). The material used for the pottery is silt and silty clay. The pots are made with a flat bottom and decorated with incisions and dots made with an obliquely held point. The patterns are geometric. The majority of the stone tools at an early stage include segments, and at a later stage these are mainly trapezes with a planed-off surface. Polished and drilled tools also occur. (Yudin, 2004).

The Early Eneolithic North Caspian culture is widespread in the North Caspian and the Volga steppes (Kurpezhe-Molla, Oroshaemoe, Algay, etc.). The pottery is made from silty clays tempered with previously fired crushed shells. Distinctive features of the North Caspian pottery are a thickened lip in the shape of a wedge and a comb pattern decoration. For stone tools quartzite was mainly used as a raw material. The main types of tools include end scrapers made from plaques and flakes, large knife-shaped blades, knives with saber-shaped and straight blades, symmetrical points and arrowheads in the shape of a “fish” (Barynkin, Vasiliev, 1985, pp. 5-73; Doga, 2023).

The Middle Eneolithic Khvalynsk culture is expressively represented by such North Caspian settlements as Kara-Khuduk, Kair-Shak VI and Kombak-Te. The pottery is made from silty clays tempered with crushed shells. The edges of the pot mouth are straight or have a molding in the shape of a projecting lip. Woven textures or a toothed stamp were used as a decorative tool, sometimes in combination with incisions or dots made by pointed implements. The tools are represented by the following categories: scrapers with retouched lateral edges, symmetrical points, straight-bladed knives, triangular arrowheads, mounted flint flakes (Barynkin et al., 1988; Barynkin, 1989; Barynkin, 2010).

It should be noted that the sites of Baibek (Grechkina et al., 2022), Kair-Shak III (Vasiliev et al., 1989), Tenteksor (Vasiliev et al., 1986), Taskuduk (Doga et al., 2023), Priozernaya (Grechkina et al., 2023) are homogeneous, that is, they do not contain any non-cultural intrusive finds or residuals from the later periods. The Varfolomeevskaya, Algay and Oroshaemoe camps are stratified, or in other words, the Neolithic and Eneolithic cultural layers are overlaid by sterile layers (Vybornov et al., 2018, Vybornov et al., 2020, Vybornov et al., 2021b; Vybornov et al., 2022b; Vybornov et al., 2020b) (Fig. 2, Photos 1-4). The Kurpezhe-Molla camp relating to the North Caspian culture has yielded an extremely insignificant number of Khvalynsk finds which date to the Middle Eneolithic period. The cultural context of the Khvalynsk sites of Kara-Khuduk (Barykin et al., 1988) and Kair-Shak VI (Barynkin, 1989) dates to the Middle Eneolithic period only. The assemblage from the Kombak-Te camp, apart from the Khvalynsk artifacts, contains a certain amount of Bronze Age finds (Barynkin, 2010).

**Fig. 2.**
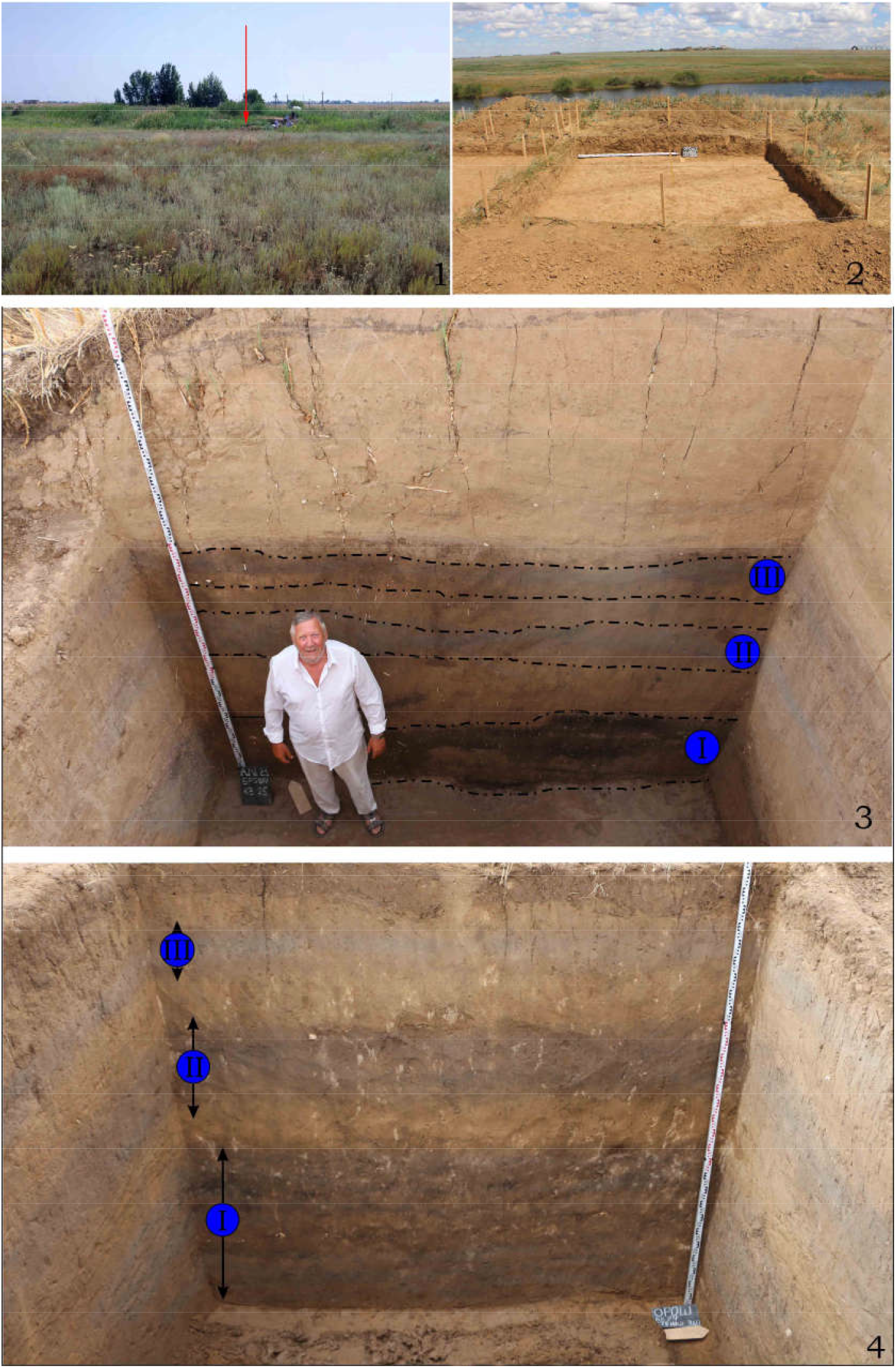
Layer stratigraphy of the excavation unit sidewall at the Algay and Oroshaemoe settlements. 1.View of the Algay settlement. 2.View of the Oroshaemoe settlement. 3.Excavation profile at the Algay settlement: I – the layer of the Late Neolithic Orlovka culture; II – the layer of the Early Eneolithic North Caspian culture; III – the layer of the Late Eneolithic Khvalynsk culture. Professor A.A. Vybornov, the excavation director. 4.The profile of the excavation unit sidewall at the Oroshaemoe settlement: I – the layer of the Late Neolithic Orlovka culture; II – the layer of the Early Eneolithic Caspian culture; III – the layer of the Late Eneolithic Khvalynsk culture.

### 3. Radiocarbon chronology

For the North Caspian Neolithic sites 112 radiocarbon dates on charcoal and animal bones were obtained, including 51 AMS results. The Kair-Shak occupation falls within the period from 7,200 to 6,900 BP (Vybornov et al., 2016; Vybornov, Kulkova 2021). The time span of the Late Neolithic Tenteksor complexes covers the period from 6700 to 6400 BP (Vybornov et al., 2016; Grechkina et al., 2023). The Jangar sites fit into the framework of 7,000-6,500 BP (Vybornov et al., 2016). The Orlovka time span is from 7,200 to 6,200 BP (Vybornov et al., 2022). The North Caspian culture occupation covers the period from 6500 to 6000 BP in the North Caspian, and from 5900 to 5800 BP in the Volga steppes (Doga, 2022). The Khvalynsk culture developed within the time span of 5900 – 5400 BP in the North Caspian region, and from 5700 to 5200 BP in the Volga steppes (Doga, 2024).

### 4. Changes in animal ratios through time

The two-millennia time span is represented by the Neolithic and subsequent Eneolithic archaeological cultures. Eight Neolithic complexes contain a total of 11,296 identifiable bones. Five Eneolithic complexes yielded 1,151 identifiable specimens. Despite such a significant difference, the number of bone samples from Eneolithic sites is quite sufficient to obtain reliable data. Nonetheless throughout the course of the Neolithic period the cultural layers of the archaeological sites yield only wild animal bones. The bones of sheep and goats first appear in the Early Eneolithic, represented by the North Caspian culture. Cattle bones appear at the Late Eneolithic Khvalynsk sites, and the number of sheep and goats also increased at that time. The hunted species are kulan, saiga, aurochs, and wild horse. All of them, with the exception of the aurochs, remained huntable throughout the period. The transformation of preferences in hunting is quite clearly represented in the diagrams. It is important to note that the percentage ratios based on the number of identifiable specimens and on the minimum numbers of individual animals do not contradict each other (Fig. 3, 4; Table 1). This confirms that deposits of animal bone remains in the cultural layers provide a reliable source. Dogs, domestic animals possibly used in hunting, have been also included into the statistical data. Apart from the game animals most frequently encountered at these settlements, the occasional bones of red deer (Cervus elaphus), wild boar (Sus scrofa), hare (Lepus europaeus), wolf (Canis lupus), fox (Vulpes vulpes), and badger (Meles meles) were found in the cultural layers, but they are relatively scarce and not present in every site.

**Fig. 3.**
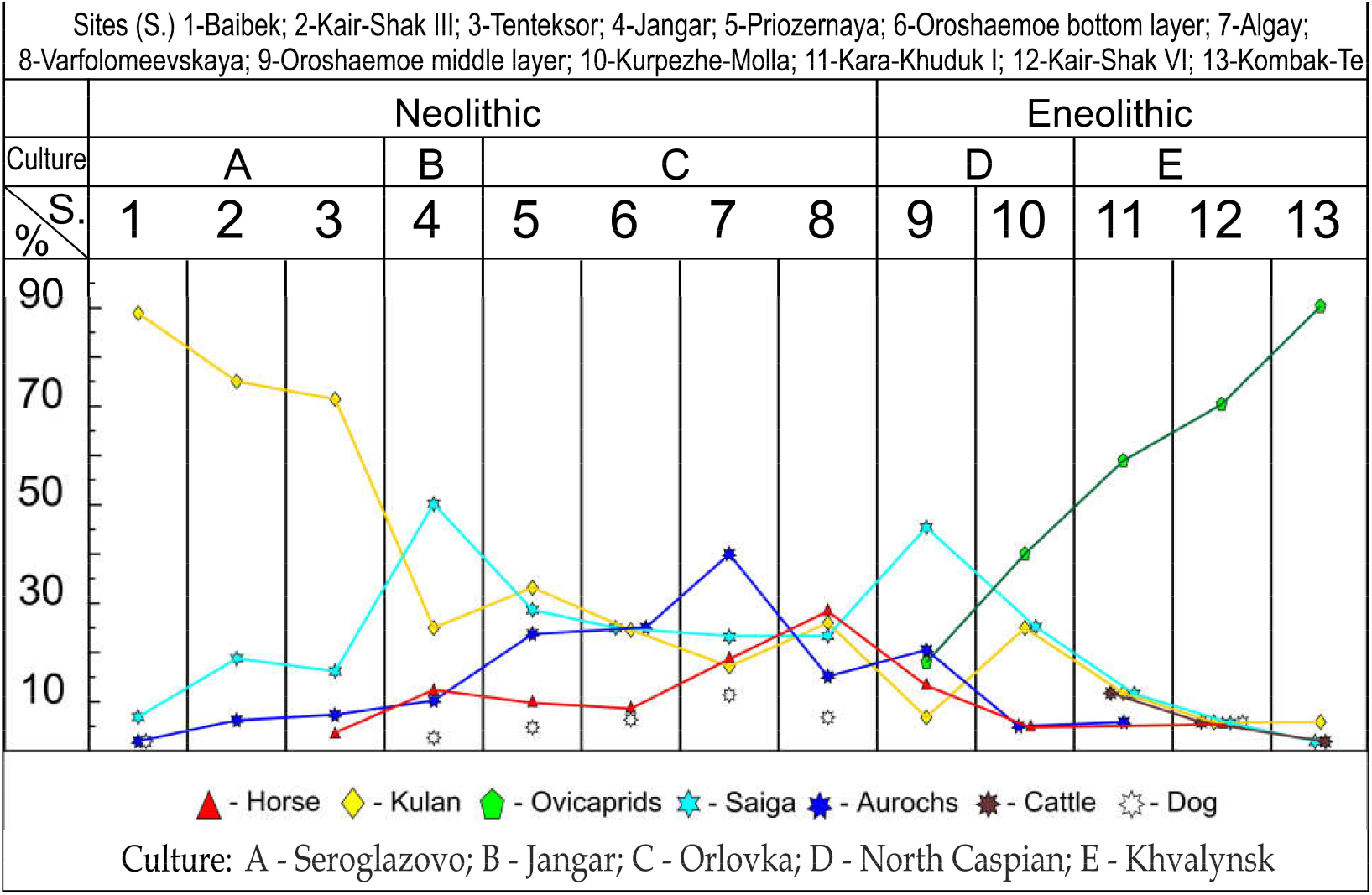
Percentage of animals based on the minimum number of individuals.

**Fig. 4.**
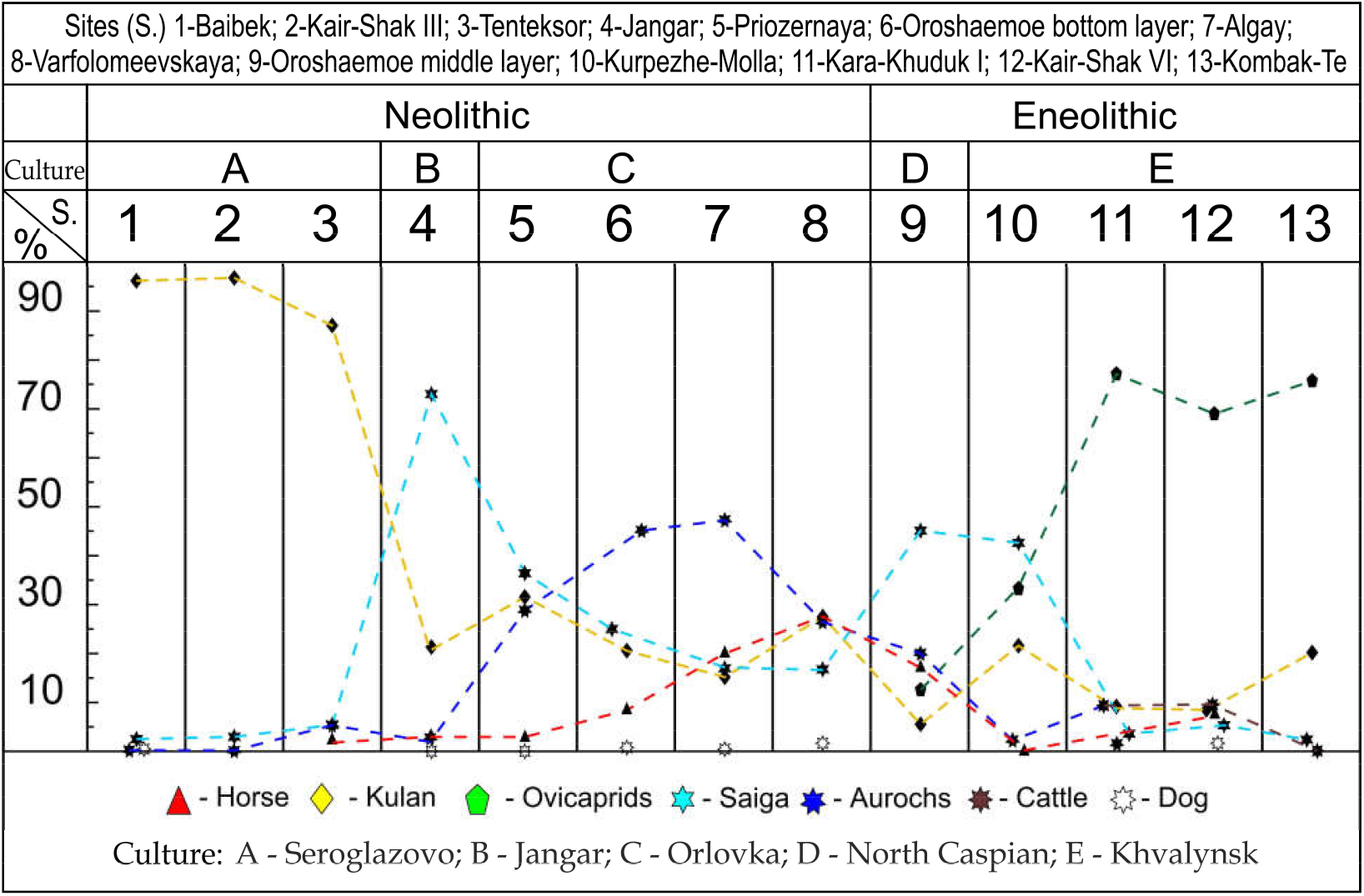
Percentage of animals based on the number of identifiable bone specimens.

**Table 1.**
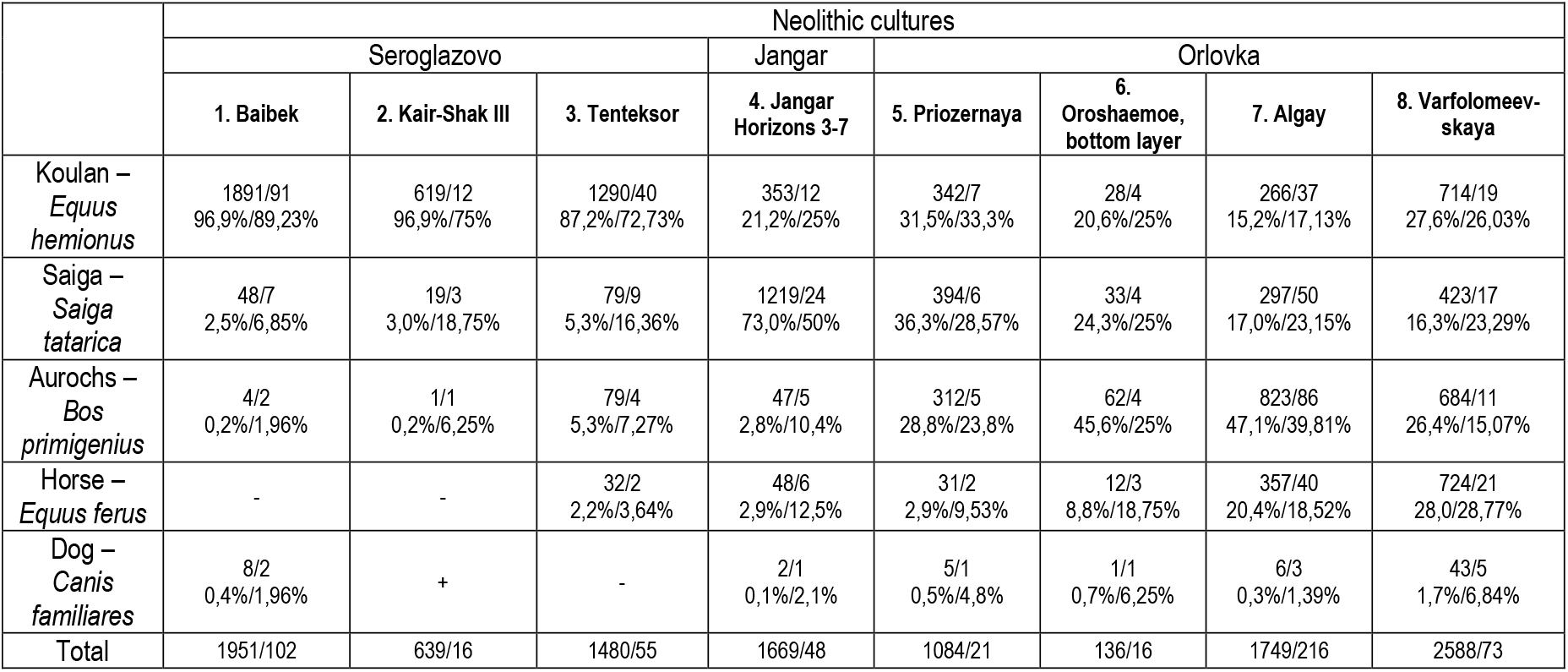
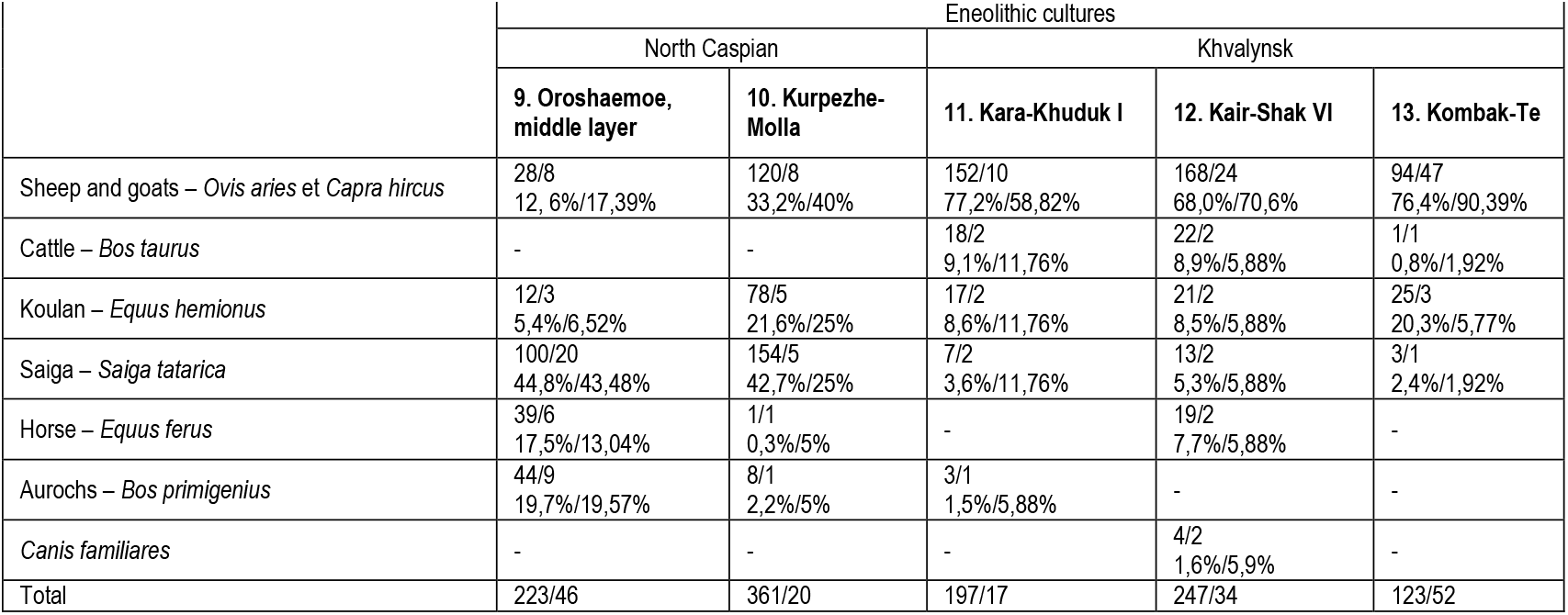
The species composition and the number of animal bone remains. The numerator indicates the number and percentage of bones. The denominator indicates the number and percentage of individual animals.

|  | Neolithic cultures |  |  |  |  |  |  |  |
| --- | --- | --- | --- | --- | --- | --- | --- | --- |
|  | Seroglazovo |  |  | Jangar | Orlovka |  |  |  |
|  | 1. Baibek | 2. Kair-Shak III | 3. Tenteksor | 4. Jangar Horizons 3-7 | 5. Priozernaya | 6. Oroshaemoe, bottom layer | 7. Algay | 8. Varfolomeevskaya |
| Koulan – *Equus hemionus* | 1891/91<br>96,9%/89,23% | 619/12<br>96,9%/75% | 1290/40<br>87,2%/72,73% | 353/12<br>21,2%/25% | 342/7<br>31,5%/33,3% | 28/4<br>20,6%/25% | 266/37<br>15,2%/17,13% | 714/19<br>27,6%/26,03% |
| Saiga – *Saiga tatarica* | 48/7<br>2,5%/6,85% | 19/3<br>3,0%/18,75% | 79/9<br>5,3%/16,36% | 1219/24<br>73,0%/50% | 394/6<br>36,3%/28,57% | 33/4<br>24,3%/25% | 297/50<br>17,0%/23,15% | 423/17<br>16,3%/23,29% |
| Aurochs – *Bos primigenius* | 4/2<br>0,2%/1,96% | 1/1<br>0,2%/6,25% | 79/4<br>5,3%/7,27% | 47/5<br>2,8%/10,4% | 312/5<br>28,8%/23,8% | 62/4<br>45,6%/25% | 823/86<br>47,1%/39,81% | 684/11<br>26,4%/15,07% |
| Horse – *Equus ferus* | - | - | 32/2<br>2,2%/3,64% | 48/6<br>2,9%/12,5% | 31/2<br>2,9%/9,53% | 12/3<br>8,8%/18,75% | 357/40<br>20,4%/18,52% | 724/21<br>28,0%/28,77% |
| Dog – *Canis familiares* | 8/2<br>0,4%/1,96% | + | - | 2/1<br>0,1%/2,1% | 5/1<br>0,5%/4,8% | 1/1<br>0,7%/6,25% | 6/3<br>0,3%/1,39% | 43/5<br>1,7%/6,84% |
| Total | 1951/102 | 639/16 | 1480/55 | 1669/48 | 1084/21 | 136/16 | 1749/216 | 2588/73 |

|  | Eneolithic cultures |  |  |  |  |
| --- | --- | --- | --- | --- | --- |
|  | North Caspian |  | Khvalynsk |  |  |
|  | 9. Oroshaemoe, middle layer | 10. Kurpezhe-Molla | 11. Kara-Khuduk I | 12. Kair-Shak VI | 13. Kombak-Te |
| Sheep and goats – *Ovis aries* et *Capra hircus* | 28/8<br>12,6%/17,39% | 120/8<br>33,2%/40% | 152/10<br>77,2%/58,82% | 168/24<br>68,0%/70,6% | 94/47<br>76,4%/90,39% |
| Cattle – *Bos taurus* | - | - | 18/2<br>9,1%/11,76% | 22/2<br>8,9%/5,88% | 1/1<br>0,8%/1,92% |
| Koulan – *Equus hemionus* | 12/3<br>5,4%/6,52% | 78/5<br>21,6%/25% | 17/2<br>8,6%/11,76% | 21/2<br>8,5%/5,88% | 25/3<br>20,3%/5,77% |
| Saiga – *Saiga tatarica* | 100/20<br>44,8%/43,48% | 154/5<br>42,7%/25% | 7/2<br>3,6%/11,76% | 13/2<br>5,3%/5,88% | 3/1<br>2,4%/1,92% |
| Horse – *Equus ferus* | 39/6<br>17,5%/13,04% | 1/1<br>0,3%/5% | - | 19/2<br>7,7%/5,88% | - |
| Aurochs – *Bos primigenius* | 44/9<br>19,7%/19,57% | 8/1<br>2,2%/5% | 3/1<br>1,5%/5,88% | - | - |
| *Canis familiares* | - | - | - | 4/2<br>1,6%/5,9% | - |
| Total | 223/46 | 361/20 | 197/17 | 247/34 | 123/52 |

Below we consider the variation through time in the game/domestic animal ratio based on each species of major importance.

#### Kulan (*Equus hemionus*)

During the Early Neolithic, represented by the Seroglazovo sites, kulan considerably outnumbers other animals. Kulan bones markedly predominate over other species. Percentages range from 96.4% to 87.2% for bones and from 89.23% to 72.73% for individual animals. Hunting is likely to have been focused on this species in the early Neolithic. The specific feature of kulans as a species is that in winter they gather in large herds of up to a hundred animals (Przhevalsky, 1888. p. 25). Probably, this influenced the choice of this species as eligible for hunting. Subsequently, the share of kulan as game gradually decreased and reached its minimum of 5.77% in the Late Eneolithic.

#### 2. Saiga (*Saiga tatarica*)

Saiga is the second largest group of hunted species after kulan. During the Late Neolithic and Early Eneolithic, the share of saiga was almost equal to that of kulan, except for the two sites: Jangar (50% of animal individuals, 73% of animal bones) and Oroshaemoe, the middle layer (48.48% of individual animals, 44.8 % of animal bones). Such a high percentage is explained by the location of these settlements on the border of the kulan distribution area. At the same time, the saiga distribution area was much larger. It covered the entire steppe zone of the northern part of Eurasia. In the Late Eneolithic, the saiga percentage decreased to 1.92% of all game animals.

#### 3. Aurochs (*Bos primigenius*)

In the Early Neolithic the aurochs percentage is relatively small. Then it begins to increase gradually. It reaches its maximum (47.01%/39.81%) in the Algay settlement in the Late Neolithic. As a result, in this settlement aurochs is predominant. Then its share as game begins to decrease, reaching its minimum in the settlements of Kurpezhe-Molla (2.2%/5%) and Kara-Khuduk (1.5%/5.88 %). In the other two Late Eneolithic settlements (Kair-Shak VI and Kombak-Te) already involved in cattle breeding, no aurochs were identified.

#### 4. Wild horse (*Equus ferus*)

Horse bones are a rare find in the Early Neolithic cultural layers. During the Late Neolithic, a gradual increase in both bones and individual animals is recorded with its maximum in the Varfolomeevskaya camp reaching 28.0%/28.77%. During the Eneolithic period, the horse percentage is significantly reduced, lacking entirely in the two Khvalynsk settlements (Kara-Khuduk and Kombak-Te). In general, hunting wild horses was not a priority. The “tarpan” as a wild species of the Volga-Ural steppes existed at least until the first half of the 19th century (Eversman, 1850. p.215-220). At that time Tarpan hunting was most successful in winter, when snow was deep (Pallas, 1773, p. 316-317; Eversman, 1850, p. 216).

It is very important to emphasize that the genetic profile of the horses inhabiting the Volga region at the time is closest to the DOM-2 profile of the first domesticated horses (Librado et al., 2021). Domesticated horses appeared much later, in the 22nd-20th centuries BCE. Their direct progenitors were found in the cultural layers of the settlements located in the Volga steppes. Their genetic profile is identified as NEO-NCAS meaning “the Neolithic of the North Caspian”. This profile was obtained from the bones of two horses found in the Varfolomeevskaya camp dating back to 57th-55th centuries BCE, one horse from the Algay camp dated to 55th-54th centuries BCE, and a horse found in the Oroshaemoe middle layer dated to 47th-45th centuries BCE. The distribution area of this genetic profile covers steppe and semi-desert zones. The distribution area of the earliest DOM-2 horses is located further north, that is in the Volga-Ural forest-steppe region. The distribution areas of domesticated and wild horses did not overlap for a long time. What prevented domesticated horses from meeting their wild kindred is a topic for separate research.

### Domesticated animals

#### 1. Dogs (*Canis familiaris*)

The dog percentage is relatively small. Some sites have no dog remains at all. Dogs are deemed to appear in the region in the Early Neolithic, approximately 6900 BP. The increase in the dog percentage corresponds to the Late Neolithic: from 0.1%/2.1% at the Jangar settlement to 1.7%/6.84% at Varfolomeevskaya site. Only the Kair-Shak VI settlement out of the five Eneolithic sites has yielded dog bones - 1.6%/5.9%.

The available ratios suggest that the increase in the dog numbers may be associated with the diversification of hunting all the game species. Groups of people living in one settlement are likely to be focused on hunting specific types of wild animals. Hence, they have specially trained dogs. Animal domestication resulted in a dramatic hunting downturn, which led to a decrease in the total number of dogs. The number of dogs required as herding or guard animals is considerably smaller compared to the needs for hunting different animals.

#### 2. Sheep and goats (*Ovis aries/Capra hircus*)

Caprines were found in the Eneolithic sites. These were either sheep or goat bones: their species are not always possible to identify accurately, though we assume that sheep as species predominated in this case. Among the identifiable bones, some certainly belong to sheep, while there is not a single one adequately identified as a goat. Sheep and goats have wild ancestors neither in the Volga nor in the Dnieper-Don region. Currently we do not know where these domestic species came from. It remains unclear whether they were brought here during direct human migration or as a result of exchange. The original distribution area where small domestic bovids originated also remains to be determined. With a high degree of caution, it is possible to assume that sheep/goats came from the territory of the Iranian Highlands moving along the eastern coast of the Caspian Sea. In this regard, it is important to draw attention to the sites located along the seashore on the Mangyshlak (or Mangghyshlaq) Peninsula, which is almost one thousand kilometers south-east away from the Lower Volga. The pottery and flint tools found in the cultural layers in these sites definitely resemble the Eneolithic finds in the Volga steppes (Astafyev, 1989).

During the Eneolithic the proportion of caprines consumed in human diet continuously increased from 12.6%/17.39% in the Oroshaemoe middle layer to 76.4%/90.39% in Kombak-Te. Over the Early Eneolithic period (the Oroshaemoe middle layer; Kurpezhe-Molla) the proportion of all game animals remains fairly large, and in the Late Eneolithic it significantly decreases.

#### 3. Cattle (*Bos taurus*)

Bones of these animals were found in each of the three Late Eneolithic settlements. But even in these settlements their share is relatively small: from 9.1% / 11.76% to 0.8%/1.92%. The bone parameters of cattle are generally not identical with those of aurochs. The cattle bones are a little smaller. This indicates that their body was more slender, which leads us to suggest that cattle were not domesticated locally, but that they were brought into the Volga region already domesticated, as well as sheep and goats. Currently it is unknown which region they came from.

## Conclusions

Over the last quarter of the 7th-5th millennia BCE, the species composition transformation of game animals constitutes a dynamic picture. In the early Neolithic, the main priority in hunting was exploiting kulan herds. In the course of the Middle and Late Neolithic, hunting was diversified, and all the species of large size (aurochs, saiga, kulan, horse) became the quarry. At the same time, the number of dog remains increases. This increase in the number of dogs we think may be connected with the fact that people from the same settlement hunted all the available species of game animals. The proportion of wild horses increased during the Late Neolithic.

In the Early Eneolithic, the first domestic animals (sheep and goats) appear. The share of saiga and aurochs remains apparently high. The kulan proportion varies, and the horse proportion noticeably decreases. In the Late Eneolithic the share of ovicaprids considerably increases while the number of all game animals decreases. Cattle also appear in the same period. The horse proportion becomes extremely small, almost vanishing. Horse bones were found in only one out of the three Khvalynsk sites: Kairshak VI 7.7%/5.88%. Thus, the ratio of the number of horse bones is in inverse proportion to the increase in the size of herds of domestic animals. This is an important fact, which contradicts the idea of horse domestication in this period. According to D. Anthony horses could have been tamed in order to go ahead and break the snow and icy crust over the feeding ground for the herds following them (Anthony, 2007. p.200).

Thus, in the course of the Eneolithic period, there is an incremental increase in the size of herds of domestic animals and the role of hunting is reduced. Caprines as well as cattle laid the foundation for the development of mobile pastoralism in the Volga-Ural region. This practice becomes the blueprint for the Yamnaya people in the early Bronze Age. The Yamnaya culture is characterized by an explicit archaeological trinity of features: kurgan burial rites, metal tools and weapons, and four-wheeled vehicles. These three features reflect the efficiency of mobile pastoralism in the course of two and a half millennia. There are burials with funerary sacrifices of animals (bones and skulls). The majority of animal deposits are sheep sacrifices, and, to a lesser extent, those of cattle. There are no horse bones in the Yamnaya burials. Neither were they found in the burials of the subsequent Catacomb and Poltavka cultures. It was only at the turn of the third/second millennia BCE that the first horse sacrifices appeared in the Volga-Ural region. All of these horses belong to the domesticated DOM-2 genetic profile.

No datasets were generated or analyzed during the current study.

## Acknowledgement

The study was supported by the Russian Science Foundation and funded as a part of project No. 24 -28-00103 “The transformation of the Late Neolithic – Eneolithic cultures in the Lower Volga: a cross-disciplinary approach”.

## Notes

https://sgspu.ru

### Competing Interest Statement

The authors have declared no competing interest.

